# DeepMHC: Deep Convolutional Neural Networks for High-performance peptide-MHC Binding Affinity Prediction

**DOI:** 10.1101/239236

**Authors:** Jianjun Hu, Zhonghao Liu

**Affiliations:** Department of Computer Science and Engineering, University of South Carolina, Columbia, SC 29201, USA.; School of Mechanical Engineering, Guizhou University, Guiyang, Guizhou,550033, China

## Abstract

Convolutional neural networks (CNN) have been shown to outperform conventional methods in DNA-protien binding specificity prediction. However, whether we can transfer this success to protien-peptide binding affinity prediction depends on appropriate design of the CNN architectue that calls for thorough understanding how to match the architecture to the problem. Here we propose DeepMHC, a deep convolutional neural network (CNN) based protein-peptide binding prediction algorithm for achieving better performance in MHC-I peptide binding affinity prediction than conventional algorithms. Our model takes only raw binding peptide sequences as input without needing any human-designed features and othe physichochemical or evolutionary information of the amino acids. Our CNN models are shown to be able to learn non-linear relationships among the amino acid positions of the peptides to achieve highly competitive performance on most of the IEDB benchmark datasets with a single model architecture and without using any consensus or composite ensemble classifier models. By systematically exploring the best CNN architecture, we identified critical design considerations in CNN architecture development for peptide-MHC binding prediction.

## Introduction

Protein-peptide interactions are critical in many biological processes. It is desirable to develop highly accurate methods to identify peptides that bind to well-known peptide-recognition domains such as SH2, SH3, PDZ, the MHC membrane receptors, and enzymes such as kinases and phosphatases. Such prediction tools can have significant medical value in terms of immunotherapy and cosmetic value and can help to illuminate the underlying mechanisms of immune responses [1]. Compared to experimental approaches for such purpose, computational prediction methods are much more efficient and cost-effective.

A large number of algorithms have been developed for protein-peptide binding prediction, either for predicting whether a peptide can bind to a receptor protein (a classification problem) or for predicting the binding affinity (a regression problem) [2, 3]. Among all the studies, computational methods for predicting peptide binding to major histocompatibility complex (MHC) class I molecules have been the most active. This is due to the availability of large scale peptide sequence data sets with experimentally determined binding affinities or binding labels as provided in e.g. the IEDB database [4, 5] and the automated benchmark platform for comparing and tracking the progress of a variety of online algorithms [3]. There are several major challenges in peptide binding prediction to MHC class I/II molecules [6, 7]. The first challenge is the huge polymorphism of MHC genes, with several thousand allelic variants in the HLA loci. So most prediction methods have to generate different models for different MHC-I receptors with allelic variations except a few pan-specific methods such as NetMHCpan and NetMHCcons [8]. The second challenge is the unbalanced peptide samples. Some allelic variants have thousands of training samples with known binding affinities or binding labels while others may only have a few hundreds or less. The third challenge is the variant preference of the peptide lengths for different allele variants. While the most common length is 9, quite a few binding peptides have lengths of 8, 10, 11, or more. The fourth challenge is the varying affinity thresholds for different allele variants of MHC-I molecules. The fifth challenge is the nonlinear high-order or distal dependencies between different amino acid positions of the binding peptides. Simple models that only consider dependencies between neighbor positions are easy to use in practice, but fail to account for the distal dependencies that are observed in the data. On the other hand, models that allow for arbitrary dependencies are prone to overfitting, requiring regularization schemes that are difficult to use in practice for non-experts. These challenges in MHC peptide binding prediction have led to a variety of tricks in term of feature engineering, machine learning architectures, and ensemble strategies to develop more effective prediction algorithms, which however also made it difficult to use such tools. Especially, the diverse expert-designed features and their contributions to the final prediction performance have barely been characterized.

## Related Work

MHC Peptide-binding prediction algorithms can be largely categorized based on their features,machine learning models, and predicton output types (qualitative or quantitative)[9]. Sequences of the peptide and the receptor, structural information of the bound complex, physichochemical properties of the amino acids, evolutionary information, word embeeding have all been exlored to develop better predictors. In terms of machine learning algorithms, support vector machines, artificial neural networks, logisitic regression, decision trees, and the most recent deep learning methods have all been applied too [10].

Among all deep learning models, deep convolutional neural networks have achieved state-of-the-art performance in a variety of fields such as computer vision, speech recognition, and natural language understanding due to their inherent capability to learn the hierarchical features required to achieve high-performance pattern recognition. Following this success, deep CNN models have been applied to a variety of bioinformatics problems, especially in DNA/RNA motif/sequence modeling and prediction including well-know algorithms such as DeepBind [11], DeeperBind [12], and DeepCpG [13]. These CNN models for DNA binding prediction usually use CNN architectures composed of 2-5 convolution/pooling layers plus one or two fully connected layers. The number of convolution filters are usually between 30-50. The DNA/RNA sequences are usually encoded using the so-called one-hot encoding, in which each nucleotide is encoded as one column vector of 4 components corresponding to A/C/G/T or A/U/G/T. The component corresponding to the nucleotide will be set as 1 and the remaining set as 0. So a DNA/RNA sequences can be represented by a two-dimension matrix. A one-dimension multi-channel convolution layer is then used to accept these inputs. To identify the best architecture for DNA-protein binding prediction, [14] conducted a systematic examination of how to design a CNN architecture for effective DNA-protein binding specificity prediction.

Successful applications of deep neural networks for protein binding prediction have also been reported but not as active as DNA binding prediction algorithms. Kuksa et al. applied a Gaussian restricted Boltzmann machine(RBM) based deep neural network (DNN) and a semi-RBMs based feed-forward high-order neural network(HONN) to MHC-peptide prediction [7]. BLOSUM encoding is used to encode peptides into 180-dimensional continuous vectors. They found that their HONN network when combined with a high-order kernel SVM method achieved better results than DNN, HONN, and NetMHC. Vang and Xie [15] applied a deep convolutional neural network model for HLA peptide-binding prediction with sophisticated word vector encoding borrowed from natural language processing. Their CNN model includes 32 convolution filters of size 7. They showed their HLA-CNN model has better performance than NetMHCpan on more than half of the benchmark datasets from IEDB. Bhattacharya et al. [16] recently also reported to use deep learning models for MHC-I binding affinity prediction. Out of all deep learning algorithms, they found that CNN based algorithms are least successful. So far, there is no report that has successfully applied deep CNN models to the MHC-I peptide binding prediction problem.

Our CNN model based approach is different from previous (Deep) neural network based approaches [17]. While artificial neural networks have been applied to MHC-peptide binding prediction [6], their performances have been limited by their feature-based encoding along with the shallow feed-forward fully connected structures instead of the convolutional neural networks as used in our approach. A major difference is that our CNN based approach takes the raw discrete amino acid sequences as input rather than expert-designed features. The simplicity of our deep CNN model for MHC-I binding affinity prediction and its superior performance are in align with the success of convolutional neural networks for DNA and RNA binding predictions[11], and methylation states prediction in CpG islands [13].

In this article, we propose a novel deep convolutional neural network based protein-peptide binding affinity prediction algorithm and demonstrated its superior prediction performance compared to the state-of-the art algorithms on a large number of peptide-binding data sets of the MHC-I class molecules. Our algorithm is also shown to perform better than other deep learning models and CNN models. Most interestingly, the overwhelming better performance of DeepMHC is achieved by a simple CNN model which only uses the raw amino acid sequences of the peptides as input without tedious, tricky, expert-based feature extraction or encoding. We don’t use any of these features such as k-mers, physicochemical properties, peptide-MHC interactions, BLOSUM, profile HMM encoding, or distributed word vector representation. Our approach used a single model structure to learn the prediction models for different allele variants of the MHC molecules. We also did not need to resort to the complex ensemble machine learning models either, which makes the training and testing complicated [6, 18].

Our contributions can be summarized in the following aspects:

- We present a systematic exploration of CNN architectures for predicting MHC-I peptide binding using large-scale public datasets. This is complementary to the similar work that explores effective way to applying CNNs to DNA-protein binding prediction [14].
- We identified a top-performing CNN model that outperforms all the traditional non-deep learning based prediction algorithms. It also works better than earlier DNN and CNN based approaches
- We conducted comprehensive comparison experiments to compare our DeepMHC algorithm to other algorithms

## Materials and Methods

### Protein Sequence Encoding

We used the one-hot encoding approach for mapping the peptide amino acid sequence into the input layer of the convolutional neural network model. As shown in Figure 1, each amino acid of the peptide is encoded by a sparse column vector of dimension 20 with corresponding component set to 1 and remaining 19 components set as 0. With the one-hot representation, there are still two different ways to map the encoded matrix to a tensor in implementation level. One is to map the input matrix as a 20-channel 1-row 2D tensor (or 20-channel 1D tensor), as is done in our proposed approach. Another way is to map the input matrix as a single channel 20-row 2D tensor. In, we compare and analyze two different encoding ways’ performance and indicate that the latter encoding way comes with a drawback.

**Figure 1.**
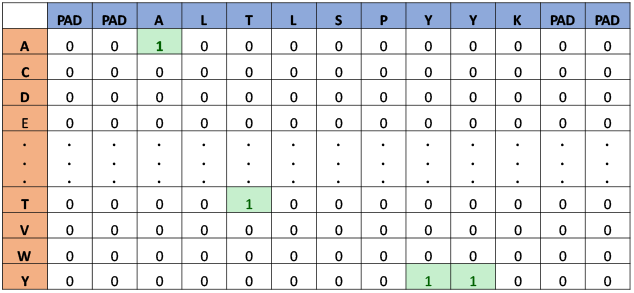
One-hot encoding of protein sequence ALTLSPYYK.

### Convolutional Neural Network Model

Out of many deep learning architectures, the convolutional neural network is among the best performing models for pattern recognition in areas such as computer vision, speech recognition, and natural language processing [19]. Our CNN based DeepMHC models are composed of two stacked convolutional layers, one max-pooling layer, and one fully connected layer (Figure 2). As shown in Figure 2, an input peptide sequence is first encoded into an 2D input tensor with shape (1,13,20). Adam [20] is used as the back-propagation optimization algorithm for training the networks.

**Figure 2.**
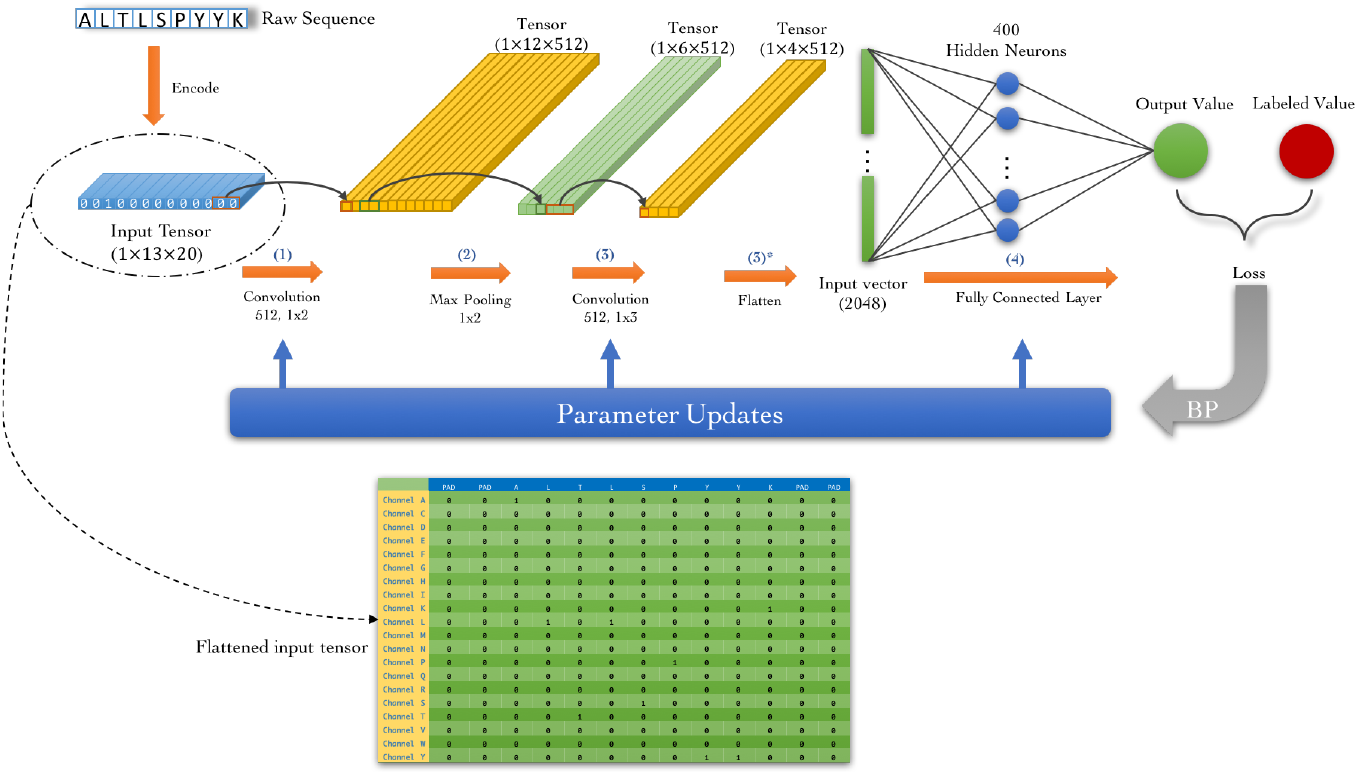
Architecture of DeepMHC Model.

#### The convolution layers

we used 512 filters of size 1x2 for the first convolutional layer and 1x3 for the second convolutional layer. Since each sequence input in our CNN model has 20 channels compared to 4 channels used in DNA nucleotide sequence encoding [11], we set the filter number to 512, which is significantly more than the 30-50 filters used in DNA binding prediction. These larger set of filters are necessary to capture the majority of the building block motifs of length 2, which can be further combined into longer motifs by succeeding layers. We found that the number of filters have dramatic effect on the performance of the CNN models as discussed later.

#### Max-pooling layers

the pooling layers are used for summarizing the activations of adjacent neurons. Different from the convolution layers in which the convolution filters move along the sequence by stride size of 1 with overlapping, the nonoverlapping pooling is applied with a stride size (1, *N*) to reduce the dimension of the input sequence and thus the number of model parameters. Two commonly used poolings are Average-Pooling and Max-Pooling. Here the max-pooling layers are used in our CNN models for summarizing the activations of adjacent *N* neurons by their maximum value. In our model, we use 1 × 2 as pooling window size.

#### Fully connected layer

As shown in Figure 2, we added one fully connected layer with the tanh activation function after flatting the output from the max-pooling layer. The fully connected layer is used to learn the non-linear mapping relationship between the extracted higher level features and the binding affinity values or binary binding labels. We used 400 hidden units in our fully connected layer and applied 50% dropout rate for regularization.

### Benchmark Data Sets and Evaluation Metrics

To ensure fair comparison with existing peptide binding prediction algorithms, we used the public IEDB training and test datasets released on the IEDB website. These datasets can be found at http://tools.iedb.org/main/datasets/. We trained on MHC-I alleles with at least about 2000 training samples from the BD2013 dataset [21]. The details of training datasets are listed in Table 1.

**Table 1.** Training Datasets from IEDB

| MHC | Data Type | Length | Seq Count | Pos | Neg | 5-fold CV Pearson |
| --- | --- | --- | --- | --- | --- | --- |
| HLA-A*02:01 | IC<sub>50</sub> | 9 | 9051 | 3273 | 5778 | 0.91 |
| HLA-A*03:01 | IC<sub>50</sub> | 9 | 5488 | 1378 | 4110 | 0.87 |
| HLA-A*11:01 | IC<sub>50</sub> | 9 | 4544 | 1363 | 3181 | 0.89 |
| HLA-A*02:03 | IC<sub>50</sub> | 9 | 4428 | 1543 | 2885 | 0.90 |
| HLA-B*15:01 | IC<sub>50</sub> | 9 | 4101 | 1187 | 2914 | 0.86 |
| HLA-A*31:01 | IC<sub>50</sub> | 9 | 3945 | 1015 | 2930 | 0.89 |
| HLA-A*01:01 | IC<sub>50</sub> | 9 | 3902 | 539 | 3363 | 0.81 |
| HLA-B*07:02 | IC<sub>50</sub> | 9 | 3868 | 1061 | 2807 | 0.80 |
| HLA-A*26:01 | IC<sub>50</sub> | 9 | 3766 | 409 | 3357 | 0.87 |
| HLA-A*02:06 | IC<sub>50</sub> | 9 | 3733 | 1522 | 2211 | 0.88 |
| HLA-A*68:02 | IC<sub>50</sub> | 9 | 3672 | 835 | 2837 | 0.90 |
| HLA-B*08:01 | IC<sub>50</sub> | 9 | 3027 | 715 | 2312 | 0.86 |
| HLA-B*58:01 | IC<sub>50</sub> | 9 | 2984 | 655 | 2329 | 0.88 |
| HLA-B*40:01 | IC<sub>50</sub> | 9 | 2824 | 478 | 2346 | 0.89 |
| HLA-B*27:05 | IC<sub>50</sub> | 9 | 2811 | 443 | 2368 | 0.92 |
| HLA-A*30:01 | IC<sub>50</sub> | 9 | 2565 | 736 | 1829 | 0.89 |
| HLA-A*69:01 | IC<sub>50</sub> | 9 | 2558 | 248 | 2310 | 0.82 |
| HLA-B*57:01 | IC<sub>50</sub> | 9 | 2529 | 384 | 2145 | 0.84 |
| HLA-B*35:01 | IC<sub>50</sub> | 9 | 2514 | 821 | 1693 | 0.92 |
| HLA-A*02:02 | IC<sub>50</sub> | 9 | 2465 | 1119 | 1346 | 0.90 |
| HLA-A*24:02 | IC<sub>50</sub> | 9 | 2395 | 496 | 1899 | 0.89 |
| HLA-B*18:01 | IC<sub>50</sub> | 9 | 2315 | 230 | 2085 | 0.81 |
| HLA-B*51:01 | IC<sub>50</sub> | 9 | 2239 | 233 | 2006 | 0.82 |
| HLA-A*29:02 | IC<sub>50</sub> | 9 | 2110 | 532 | 1578 | 0.86 |
| HLA-A*68:01 | IC<sub>50</sub> | 9 | 2036 | 830 | 1206 | 0.90 |
| HLA-A*33:01 | IC<sub>50</sub> | 9 | 1929 | 387 | 1542 | 0.82 |
| HLA-A*23:01 | IC<sub>50</sub> | 9 | 1915 | 410 | 1505 | 0.86 |
| HLA-A*02:01 | IC<sub>50</sub> | 10 | 2753 | 1150 | 1603 | 0.79 |
| HLA-A*03:01 | IC<sub>50</sub> | 10 | 1694 | 720 | 974 | 0.67 |
| HLA-A*11:01 | IC<sub>50</sub> | 10 | 1680 | 755 | 925 | 0.79 |
| HLA-A*68:01 | IC<sub>50</sub> | 10 | 1651 | 740 | 911 | 0.73 |
| HLA-A*31:01 | IC<sub>50</sub> | 10 | 1640 | 576 | 1064 | 0.65 |
| HLA-A*02:06 | IC<sub>50</sub> | 10 | 1623 | 679 | 944 | 0.52 |
| HLA-A*68:02 | IC<sub>50</sub> | 10 | 1617 | 481 | 1136 | 0.68 |
| HLA-A*02:03 | IC<sub>50</sub> | 10 | 1614 | 811 | 803 | 0.51 |
| HLA-A*33:01 | IC<sub>50</sub> | 10 | 1560 | 315 | 1245 | 0.48 |
| HLA-A*02:02 | IC<sub>50</sub> | 10 | 1445 | 658 | 787 | 0.73 |

The evaluation dataset were downloaded from IEDB’s weekly benchmark dataset [10]. We downloaded all datasets from 2014-03-21 to 2016-12-09 with test samples combined according to alleles, sequence length and measure type. We separated the evaluation datasets into two groups: one with affinity scores: *IC*_50_ and t1/2 (Table 2) and another with binary affinity labels (Table 3).

**Table 2.**
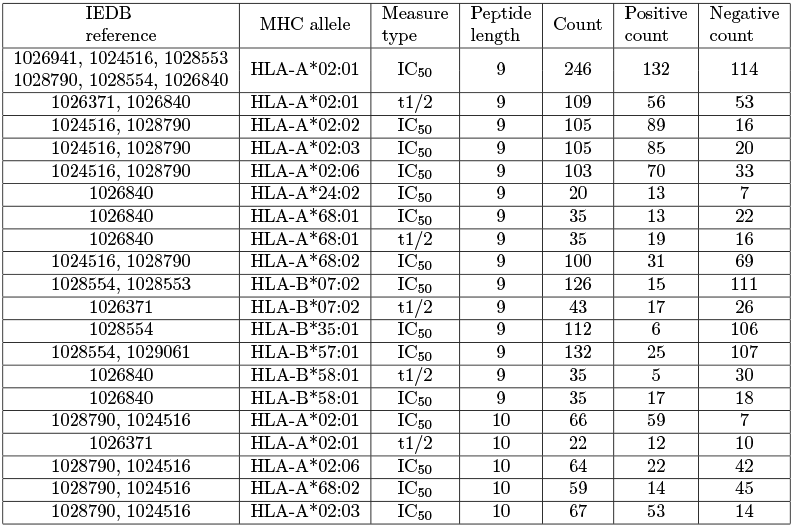
Test datasets with affinity scores

**Table 3.**
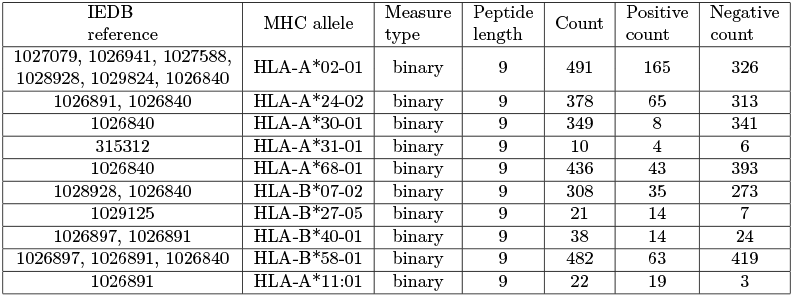
Test datasets with binary affinity labels

Following the metrics used in [10],area under the receiver operating curve (AUC) and the Spearman rank correlation coefficient (SRCC), and Pearson coefficient (Pearson) were used as metrics. A special attention is needed when calculating metrics involving with *IC*_50_: the labeled and predicted *IC*_50_ values should be negatived since the higher *IC*_50_ represents low binding preference.

## Results

### Evaluation on Benchmark Dataset

To compare with other MHC-I peptide binding prediction methods evaluatd with IEDB’s benchmark dataset, our CNN models were trained on the BD2013 dataset. The testing data were extracted from IEDB’s MHC-I weekly testing data. Both datasets are described in detail in Sections Table 1 and Table 2. Our CNN network structure is presented in Figure 2 and we used Mean Squared Error (MSE) as the loss function.

The peptide binding affinity values in BD2013 are measured in IC_50_ distributed in a large real value range (0.0 ~ 80000.0), which makes it difficult for the neural network models to predict. So we converted IC_50_ to pIC_50_ by:

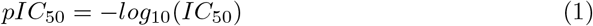

In this way, the CNN network could conduct effective search in a much smaller real value range. In all the following experiments, we used pIC_50_ as the default affinity values for training our CNN models except explicit declaration of other cases.

Five-fold cross-validation was applied to the samples for each MHC-I allele in Table 1. We first split the samples into 5 subsets of equal size and then each of the subset is tested on the model trained with other 4 subsets following this procedure:

#### Training

First, we configure a CNN model described in section and train this model for up to 5000 epochs. Before each training epoch, the training samples are randomly split into training and validation sets following 9:1 ratio. The training stops if the minimum validation loss haven’t reduced for 100 epochs (known as early stop) or reached 5000 epochs. We trained models for each allele listed in Table 1 and we trained separate models for the same allele with different peptide lengths.

#### Evaluation results of cross-validation

For sequences in each subset held out as the test set, we applied the trained CNN model to predict their affinity values. This process is repeated 5 times for all 5 subsets. And then the final Pearson correlation coefficients of all peptide samples of a MHC-I allele are calculated. The 5-fold cross-validation performances of the CNN models on all MHC-I alleles are listed in Table 1. The Pearson scores range from 0.48 to 0.92. Our CNN models achieved Pearson scores of more than 0.8 for 27 out of the 37 alleles. We also found that most of the low-performance scores are from the alleles with fewer number of peptide sequences (i 1000), which indicates that sufficient number of peptide sequences are required for our CNN models to achieve good performance. It is also found that all these low-performance allele dataset consist of peptide sequences of length 10.

#### Evaluation results on test datasets

For each peptide sequence in the testing dataset in Table 2, we applied the 5 trained cross-validation models for each allele and predict 5 affinity values. And then the average of these predicted values from all 5 models is set as the final predicted *pIC*_50_ for the peptide sequence. Finally, the predicted *pIC*_50_ values are converted back into *IC*_50_ values based on Equation 1. And then the corresponding AUC, SRCC and Pearson scores are calculated accordingly, which are then compared to the prediction results of other algorithms included in IEDB’s weekly benchmark report. The compared algorithms are: NetMHCpan [22], SMM [23], ANN [24], ARB [25], NetMHCcons [26], SMMPMBEC [27], IEDBconsensus [28] and PickPocket [29]. The complete predicted binding affinities of our method on benchmark dataset can be found in the supporting materials.

### Performance of DeepMHC on MHC-I Binding Affinity Prediction

Table 4 shows the performance comparison results on 15 datasets of peptides with length 9 for DeepMHC and 7 other prediction algorithms. Three evaluation criteria are used here including AUC, SRCC, and Pearson Coefficient. First, we highlight the best AUC/SRCC/Pearson scores for each dataset and count the number of datasets that each algorithm achieved the best scores. In the second to last row of Table 4, we found that DeepMHC has achieved best performance on 6/7/6 datasets out of 15 in terms of AUC/SRCC/Pearson respectively, which is significantly better than the best existing algorithm NetMHCpan, which achieved best performance on 3/5/2 datasets out of 15 in terms of AUC/SRCC/Pearson. It also outperformed the second best existing algorithm, ANN, which achieved best performance on 4/2/4 datasets out of 15 in terms of AUC/SRCC/Pearson. It is found that all other 4 algorithms only achieved best performance on one or two datasets. We also found that our models didn’t obtain any best results with three metrics in only 6 test allele datasets. We further calculated the average performance scores across all 15 datasets of all compared algorithms. The DeepMHC is still the best, which indicates that our proposed convolution network models has the best overall peptide binding prediction capability across different alleles.

**Table 4.**
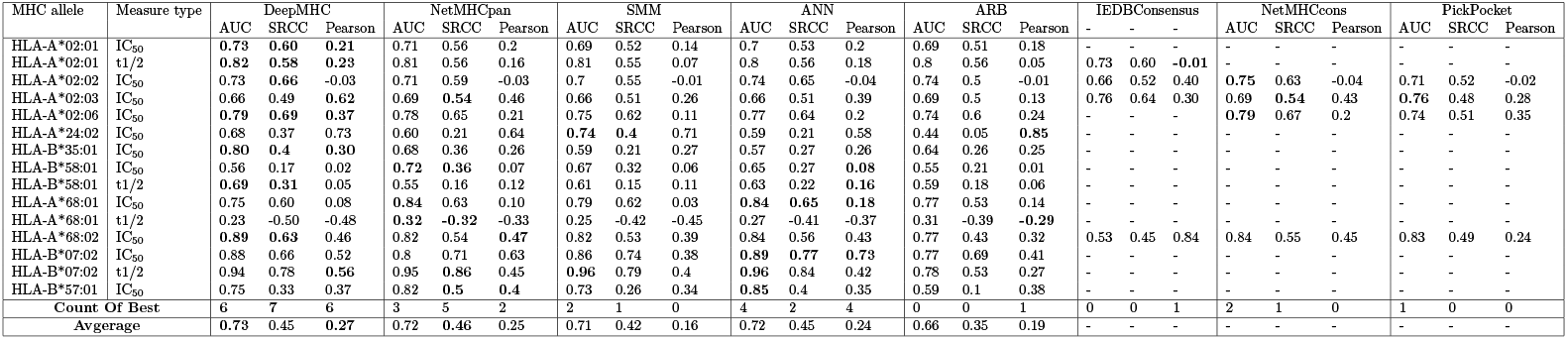
Performance of DeepMHC on binding affinity Prediction (IC_50_) compared to other algorithms (9-length, methods without winning on any dataset are omitted)

Further analysis is conducted to check the alleles that our algorithm did not achieved the best results. First, we found that for 4 test alleles [HLA-A*02:02 (IC_50_), HLA-A*24:02 (IC_50_) and HLA-B*07:02 (IC_50_ and t1/2)] on which our model didn’t get best results in terms of the AUC score, the performance of our algorithm is either very close to the best scores or among the top 3 scores.

We then analyzed the prediction results on the 5 alleles [HLA-A*02:03 (IC_50_), HLA-B*58:01 (IC_50_), HLA-B*68:01 (IC_50_ and t1/2), HLA-B*57:01(IC_50_)] that our model achieved more inferior performance compared to the best scores, trying to figure out possible reasons.

1. **Sample Size** As shown in Table 1, the HLA-B*68:01 allele dataset has only 2036 training samples, which may have caused the lower performance of DeepMHC. It is well known that the number of training samples should be large enough for DNN models to achieve good prediction performance.
2. **Mislabelled test samples in some datasets** In the training dataset BD2013, IEDB combines KD (thermo-dynamic constant), IC_50_ and EC_50_ together represented as IC_50_ since IC_50_ and EC_50_ measurements can approximate KD [10]. And we found that in the IEDB weekly benchmark data, some test peptide sequences also appear in the training data, but labelled with different binding affinities due to different measuring techniques. One example is the ELAAHQKKI peptide, which binds to allele HLA-A*02:03. In the training data, its affinty value is labeled as 2650 (radioactivity dissociation constant KD). In the weekly testing data, it is labeled as value 1.0 (radioactivity half maximal inhibitory concentratio IC50). In our testing experiments, we found that our method and other benchmark methods all failed on predicting this sequence’s IC_50_ value. Since we followed the same method in [10] marking a sequence sample as negative if its IC_50_ ¡ 500.0 nm, this kind of samples would drag the AUC scores down for all algorithms. For the test entry HLA-A*02:03 (IC_50_), our model gave the best prediction performance with a Pearson score of 0.62 and low AUC score. We found among 105 testing sequences, 14 sequences appeared in the training data with conflicting affinity values. This may explain the contradictary cases where our algorithm achieved highest Pearson scores while showing worse performance in terms of AUC.

There are other possible issues that may lead to the low performance of our algorithm in some datasets, which are further explored in Section and Section along with discussion on how to overcome these issues.

It should be noted that the state-of-the-art NetMHCpan algorithm is a very time-consuming and sophisticated algorithm. It utilized a pan-specific strategy, which trained each allele’s samples on a set of artifical neural networks with 22 to 86 hidden neurons with 3 types of input encodings and then picked the best 15 ANN networks to compose an ensemble ANN prediction model[22]. Comparing with NetMHCpan, our CNN algorithm trains allele-specific models with a single fixed convolutional neural network structure and a simple one-hot encoding while performing better on more than half of the test datasets in terms of AUC scores.

Table 5 shows the performance results over the datasets with peptide sequences of 10 amino acids. It can be found that our model performed much better than all other compared algorithms with the best AUC scores in 4 out of 5 datasets, best SRCC scores for all 5 datasets, and best Pearson scores in 2 out of 5 datasets. And on the test entries HLA-A*02-01(t1/2), HLA-A*02-03(IC_50_), and HLA-A*68-02(IC_50_), our algorithm has large leading margin in terms of the accuracy scores. We also found that the alleles in Table 5 over which the DeepMHC obtained almost all the best results correspond to the same alleles in Table 4 over which the DeepMHC also performed well. This indicated that our algorithm could effectively capture the essential features for high-quality affinity prediction regardless of the variation of peptide lengths.

**Table 5.** Prediction Performance of DeepMHC on the binding affinity (IC_50_) problems against other algorithms (10-length, Omitted some methods having no best reults)

| MHC allele | Measure type | DeepMHC |  |  | NetMHCpan |  |  | SMM |  |  | SMMPMBEC |  |  | IEDBConsensus |  |  |
| --- | --- | --- | --- | --- | --- | --- | --- | --- | --- | --- | --- | --- | --- | --- | --- | --- |
|  |  | AUC | SRCC | Pearson | AUC | SRCC | Pearson | AUC | SRCC | Pearson | AUC | SRCC | Pearson | AUC | SRCC | Pearson |
| HLA-A*02-01 | t1/2 | **0.76** | **0.4** | **0.07** | 0.57 | 0.15 | -0.13 | 0.56 | 0.14 | -0.10 | - | - | - | - | - | - |
| HLA-A*02-01 | IC<sub>50</sub> | **0.69** | **0.58** | 0.09 | 0.68 | 0.46 | 0.09 | 0.66 | 0.50 | **0.16** | 0.62 | 0.5 | 0.15 | 0.66 | 0.50 | 0.09 |
| HLA-A*02-03 | IC<sub>50</sub> | **0.80** | **0.39** | -0.05 | 0.75 | 0.25 | -0.06 | 0.73 | 0.32 | 0.00 | 0.7 | 0.31 | -0.01 | 0.73 | 0.30 | **0.02** |
| HLA-A*02-06 | IC<sub>50</sub> | 0.82 | **0.66** | 0.14 | 0.81 | 0.64 | 0.15 | 0.83 | 0.64 | 0.21 | **0.84** | **0.66** | **0.22** | - | - | - |
| HLA-A*68-02 | IC<sub>50</sub> | **0.94** | **0.78** | **0.51** | 0.66 | 0.31 | 0.12 | 0.75 | 0.41 | 0.13 | 0.73 | 0.38 | 0.11 | 0.73 | 0.40 | 0.25 |
| Count Of Best |  | 4 | 5 | 1 | 0 | 0 | 0 | 0 | 0 | 1 | 1 | 1 | 1 | 0 | 0 | 0 |
| Average |  | **0.80** | **0.56** | **0.15** | 0.69 | 0.36 | 0.04 | 0.71 | 0.40 | 0.08 | - | - | - | - | - | - |

### Performance of DeepMHC on MHC-I Binary Peptide Binding Prediction

We evaluated prediction performance of DeepMHC on benchmark datasets with binary labels following the same procedure as described before and the prediction performances are shown in Table 6. First, we found that our raw amino acid sequence based DeepMHC models are competitive compared to other algorithms that depend on additional physichochemical information and sophisticated feature engineering. Out of 10 allele datasets, DeepMHC achieved top performance on 4 datasets, which is better than NetMHCpan, SMM and ANN methods. Out of the remaining 6 datasets, DeepMHC obtained competitive results on 4 datasets (HLA-A*02-01, HLA-A*68-01,HLA-B*07-02, HLA-A*31-01) with AUC differences of less than or equal to 0.04 compared to the top scores of other algorithms. But for cases like HLA-B*27:05 and HLA-A*24:02, our performance is much lower, which can be partially attributed to the reasons as mentioned in Section. To further explore the reasons, we plotted the ROC curves of DeepMHC compared to NetMHCpan on these two datasets in Figure 3. It is found that our models do not perform as well as NetMHCpan does when the false positive rate is less than 0.2.

**Figure 3.**
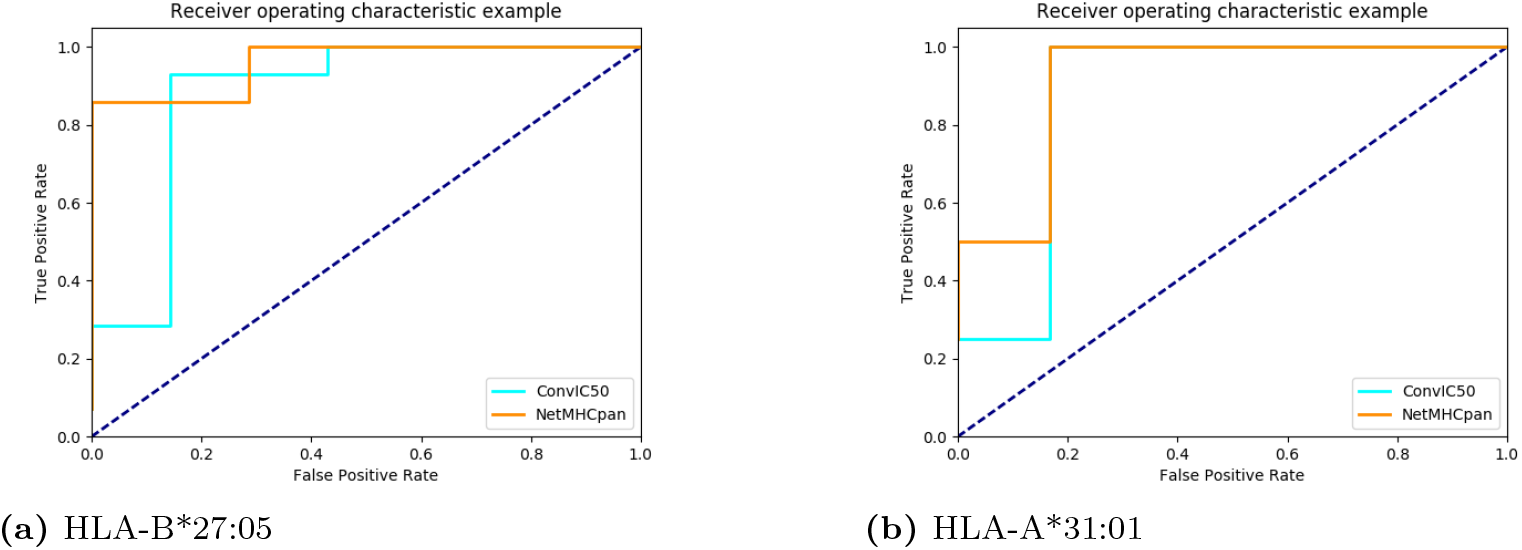
Comparing ROC curves of DeepMHC v.s. NEetMHCpan. (a) over HLA-B*27:05 dataset and (b) over HLA-A*24:02 dataset

**Table 6.** Prediction Performance of DeepMHC on binary peptide binding prediction

| MHC allele | DeepMHC |  |  | NetMHCpan |  |  | SMM |  |  | ANN |  |  |
| --- | --- | --- | --- | --- | --- | --- | --- | --- | --- | --- | --- | --- |
|  | AUC | SRCC | Pearson | AUC | SRCC | Pearson | AUC | SRCC | Pearson | AUC | SRCC | Pearson |
| HLA-A*02-01 | 0.87 | 0.60 | 0.59 | **0.89** | **0.63** | **0.66** | 0.88 | 0.62 | 0.14 | 0.88 | 0.61 | 0.64 |
| HLA-A*11-01 | **0.63** | **0.16** | **0.53** | 0.58 | 0.09 | 0.49 | 0.60 | 0.11 | 0.37 | 0.61 | 0.14 | 0.4 |
| HLA-A*24-02 | 0.81 | 0.41 | 0.24 | **0.89** | **0.51** | 0.63 | 0.87 | 0.48 | 0.23 | 0.89 | 0.51 | **0.65** |
| HLA-A*30-01 | **0.89** | **0.20** | 0.15 | 0.80 | 0.16 | 0.16 | 0.79 | 0.15 | 0.06 | 0.77 | 0.14 | **0.17** |
| HLA-A*31-01 | 0.88 | 0.64 | 0.55 | **0.92** | **0.71** | 0.74 | **0.92** | **0.71** | 0.34 | **0.92** | **0.71** | 0.7 |
| HLA-A*68-01 | 0.85 | 0.36 | 0.27 | **0.87** | **0.38** | 0.47 | 0.86 | 0.37 | 0.11 | 0.87 | 0.38 | **0.52** |
| HLA-B*07-02 | 0.90 | 0.44 | 0.42 | 0.92 | 0.46 | 0.57 | **0.93** | **0.47** | 0.09 | 0.93 | 0.47 | **0.71** |
| HLA-B*27-05 | 0.88 | 0.62 | 0.70 | **0.96** | **0.75** | **0.79** | 0.91 | 0.67 | 0.69 | 0.94 | 0.72 | 0.78 |
| HLA-B*40-01 | **0.89** | 0.65 | 0.35 | **0.89** | **0.66** | 0.42 | 0.88 | 0.63 | 0.26 | 0.88 | 0.64 | **0.48** |
| HLA-B*58-01 | **0.89** | **0.45** | 0.39 | 0.88 | 0.44 | 0.53 | **0.89** | **0.45** | 0.08 | 0.87 | 0.44 | **0.59** |
| Count of Best | 4 | 3 | 1 | **6** | **6** | 2 | 3 | 3 | 0 | 1 | 1 | **6** |
| Average | 0.85 | 0.45 | 0.42 | **0.86** | **0.48** | 0.55 | 0.85 | 0.47 | 0.24 | **0.86** | **0.48** | **0.56** |

### Comparison with Other DNN and CNN based approaches

We compared the performance of DeepMHC with three other deep neural network based algorithms including HONN [7], HLA-CNN [15], and MHCnuggets [16].

### HLA-CNN

HLA-CNN [15] is a CNN based algorithm for HLA-binding prediction using a special word-vector encoding borrowed from natural language processing. It takes protein sequences as sentences in a text corpus. Specifically individual peptides are treated as individual sentences and amino acids are treated as words. Then a skip-gram model is used with a context window of size 5 to embed each amino acid into a 15-dimensional real-value vector space. The initial embedding is learned from the set of all peptides across all MHC alleles in the training set. The output encoding of a peptide is a 2-dimensional matrix of size 9 × 15, where 9 is the length of HLA binding peptides. Their model then uses two convolution layers, each composed of 32 1D filters of size 7 together with LeakyRELU and Dropout for regularization. In their original paper, HLA-CNN was trained on a dataset assembled from several sources (IEDB, MHCBN and SYFPEITHI) and was only evaluated on 3 benchmark datasets (2015-08-7, 2016-05-03 and 2016-02-19). To ensure fair comparison, we downloaded HLA-CNN source code (via github repo: https://github.com/uci-cbcl/HLA-bind/, commit: 20ed937) provided by the author and trained HLA-CNN on BD2013 dataset. Then, we evaluated their models on the same benchmark dataset as shown in Section. The comparison results are shown in Table 7.

**Table 7.** Prediction Performance Comparison Between DeepMHC and HLA-CNN

| Allele | Measure Type | Peptide Length | Peptide Count | HLA-CNN* |  | DeepMHC |  | HLA-CNN |  |
| --- | --- | --- | --- | --- | --- | --- | --- | --- | --- |
|  |  |  |  | AUC | SRCC | AUC | SRCC | AUC | SRCC |
| HLA-A*68-01 | 9 | ic50 | 35 | 0.594 | 0.158 | **0.748** | **0.605** | 0.647 | 0.246 |
| HLA-A*24-02 | 9 | ic50 | 20 | 0.385 | -0.195 | **0.681** | **0.366** | 0.438 | -0.106 |
| HLA-A*02-02 | 9 | ic50 | 105 | **0.744** | 0.303 | 0.73 | **0.665** | 0.756 | 0.318 |
| HLA-B*35-01 | 9 | ic50 | 112 | 0.736 | 0.184 | **0.799** | **0.401** | 0.805 | 0.238 |
| HLA-B*57-01 | 9 | ic50 | 132 | **0.833** | **0.445** | 0.75 | 0.33 | 0.821 | 0.429 |
| HLA-B*58-01 | 9 | ic50 | 35 | **0.631** | **0.226** | 0.562 | 0.173 | 0.582 | 0.142 |
| HLA-A*68-02 | 9 | ic50 | 100 | **0.899** | **0.639** | 0.889 | 0.627 | 0.9 | 0.642 |
| HLA-A*02-06 | 9 | ic50 | 103 | **0.880** | 0.614 | 0.791 | **0.695** | 0.862 | 0.586 |
| HLA-B*07-02 | 9 | ic50 | 126 | 0.808 | 0.346 | **0.876** | **0.665** | 0.856 | 0.399 |
| HLA-A*02-03 | 9 | ic50 | 105 | **0.721** | 0.3 | 0.661 | **0.489** | 0.695 | 0.266 |
| HLA-A*02-01 | 9 | ic50 | 246 | **0.737** | 0.41 | 0.734 | **0.600** | 0.721 | 0.382 |
| HLA-A*11-01 | 9 | binary | 22 | **0.667** | **0.198** | 0.632 | 0.157 | 0.579 | 0.094 |
| HLA-B*27-05 | 9 | binary | 21 | 0.837 | 0.551 | **0.878** | **0.617** | 0.908 | 0.667 |
| HLA-A*31-01 | 9 | binary | 10 | **0.917** | **0.711** | 0.875 | 0.64 | 1 | 0.853 |
| HLA-B*40-01 | 9 | binary | 38 | **0.905** | **0.677** | 0.887 | 0.647 | 0.893 | 0.657 |
| HLA-A*02-01 | 9 | binary | 491 | **0.856** | **0.583** | 0.679 | 0.431 | 0.86 | 0.59 |
| HLA-A*24-02 | 9 | binary | 378 | 0.741 | 0.315 | **0.815** | **0.411** | 0.817 | 0.414 |
| HLA-B*58-01 | 9 | binary | 482 | 0.851 | 0.41 | **0.888** | **0.453** | 0.866 | 0.427 |
| HLA-B*07-02 | 9 | binary | 308 | 0.875 | 0.413 | **0.898** | **0.438** | 0.88 | 0.418 |
| HLA-A*68-01 | 9 | binary | 436 | 0.811 | 0.321 | **0.85** | **0.361** | 0.798 | 0.308 |
| HLA-A*30-01 | 9 | binary | 349 | 0.513 | 0.007 | **0.889** | **0.202** | 0.672 | 0.089 |
| HLA-A*02-01 | 10 | ic50 | 66 | 0.557 | 0.061 | **0.69** | **0.58** | 0.598 | 0.105 |
| HLA-A*02-03 | 10 | ic50 | 67 | **0.827** | **0.461** | 0.803 | 0.389 | 0.827 | 0.461 |
| HLA-A*02-06 | 10 | ic50 | 64 | **0.85** | 0.575 | 0.817 | **0.657** | 0.865 | 0.6 |
| HLA-A*68-02 | 10 | ic50 | 59 | 0.898 | 0.587 | **0.943** | **0.784** | 0.9 | 0.59 |
| Average |  |  |  | 0.763 | 0.372 | **0.791** | **0.495** | 0.782 | 0.393 |
| Count of Best |  |  |  | 13 | 8 | 12 | 17 | - | - |

During our testing we found an error in HLA-CNN’s code: it uses the testing dataset as validation set during training. Since the validation loss was used to decide when to stop training, this mistake would have led to overestimated prediction performance. To ensure a fair comparison, we trained two models for each allele with two versions of the HLA-CNN code: HLA-CNN and HLA-CNN* as shown in Table 7. Prediction results of the models trained with the original code are denoted as HLA-CNN. For HLA-CNN*, we modified the original code so that during training the CNN model would be validated only on part of the training dataset instead of using the testing data. The bug-fixed code can be found at github (https://github.com/pcpLiu/HLA-bind). Actually we just modified one line of the code.

First, we compared the performance of HLA-CNN and HLA-CNN* over 25 MHC-I allele datasets as shown in Table 7. We found that HLA-CNN obtained higher AUC scores and higher SRCC scores over 16 datasets out of 25 testing entries comparing with HLA-CNN*. This verified that the reported performance based on the original training procedure of HLA-CNN is over-estimated.

When comparing HLA-CNN* with our method, two methods obtained similar performance in terms of the AUC scores, where HLA-CNN* obtained better results on 13 testing entries and DeepMHC better on 12 entries. Among 13 entries where HLA-CNN* obtained higher AUC scores, in only 1 entry (HLA-A*02:01, 9-length, binary) its AUC score has a larger leading margin of 0.177. For all other cases, its AUC scores have small leading margins (< 0.1). Out of the 12 testing entries where our method obtained higher AUC scores, there are 4 entries that our method has much larger leading AUC score margins : 0.157 (HLA-A*68:01, 9-length, IC_50_), 0.296 (HLA-A*24:02, 9-length, IC_50_), 0.376 (HLA-A*30:01, 9-length, binary) and 0.133 (HLA-A*02:01, 10-length, IC_50_). As the result, our method obtained a higher average AUC score (0.791) than HLA-CNN* (0.763) though HLA-CNN* is designed as a binary classifier. Actually even when compared with the unmodified HLA-CNN with an average AUC score of 0.782, our method still have a higher average AUC score as shown in Table 7. In terms of SRCC, our model performed much better. HLA-CNN* obtained 8 better SRCC scores and our algorithm obtained 17 ones. This is partially due to the fact that HLA-CNN* is trained as a binary classifier. For the 8 testing entries HLA-CNN* having higher SRCC scores, there are 2 entries in which it has large leading margins (> 0.1). And out of 17 testing entries that our algorithm obtained higher SRCC scores, there are 10 entries that DeepMHC has large leading margins (> 0.1). Thus, the average SRCC score of our method (0.495) is significantly better than those of HLA-CNN* (0.372) and HLA-CNN (0.393).

### MHCnuggets

Bhattacharya et al. [16] evaluated a set of deep learning algorithms for MHC-I binding prediction including a fully connected DNN, two recurrent neural network models, and two convolutional neural network models. Their MHCnuggets-Chunky-CNN is a 1D convolutional neural network, with kernel sizes (2,3), stride 1, a global max pooling layer, a fully connected layer with 64 hidden units, and a dropout probability of 0.6, trained for 100 epochs. Their MHCnuggets-Spanny-CNN is a 1D convolutional neural network, with kernel sizes (2,3,9), stride=1, number of filters =(1,1,1), a global max pooling layer, a fully connected layer with 64 hidden units, and a dropout probability of 0.6, trained for 100 epochs.

We downloaded the code of MHCnuggets and compared their performance on the same weekly dataset. Since the downloaded code contains trained models, we directly run MHCnuggets to make the predictions on the test dataset. We retrained our models on the training data used in MHCnuggets and then tested it on the test dataset. Table 8 shows the results of both methods. We found that these two algorithms showed similar performance, which is understandable as these two models are more similar to each other in terms of input encoding and architecture.

**Table 8.** Prediction Performance of DeepMHC V.S. MHCnuggets CNN model

| Allele | Length | Measure Type | MHCnuggets-CNN |  |  | DeepMHC |  |  |
| --- | --- | --- | --- | --- | --- | --- | --- | --- |
|  |  |  | AUC | SRCC | Pearson | AUC | SRCC | Pearson |
| HLA-A*02:01 | 9 | IC<sub>50</sub> | **0.75** | **0.66** | **0.28** | 0.73 | 0.61 | 0.17 |
| HLA-A*02:01 | 9 | binary | 0.80 | 0.49 | **0.49** | **0.86** | **0.59** | 0.44 |
| HLA-A*02:01 | 9 | t1/2 | 0.73 | 0.46 | **0.27** | **0.76** | **0.52** | 0.23 |
| HLA-A*02:02 | 9 | IC<sub>50</sub> | 0.75 | **0.69** | -0.04 | **0.79** | 0.67 | **-0.02** |
| HLA-A*02:03 | 9 | IC<sub>50</sub> | 0.68 | **0.54** | 0.70 | **0.69** | 0.50 | **0.81** |
| HLA-A*02:06 | 9 | IC<sub>50</sub> | **0.86** | **0.77** | **0.42** | 0.84 | 0.74 | 0.30 |
| HLA-A*24:02 | 9 | IC<sub>50</sub> | **0.58** | **0.32** | 0.43 | 0.53 | 0.10 | **0.47** |
| HLA-A*24:02 | 9 | Binary | 0.75 | 0.32 | 0.38 | **0.78** | **0.36** | **0.40** |
| HLA-A*68:01 | 9 | IC<sub>50</sub> | 0.58 | 0.22 | **0.14** | **0.74** | **0.55** | 0.08 |
| HLA-A*68:01 | 9 | Binary | 0.74 | 0.25 | **0.26** | **0.81** | **0.32** | 0.18 |
| HLA-A*68:01 | 9 | t1/2 | **0.34** | **-0.39** | **-0.33** | 0.24 | -0.44 | -0.37 |
| HLA-A*68:02 | 9 | IC<sub>50</sub> | **0.92** | **0.77** | **0.61** | 0.86 | 0.69 | 0.55 |
| HLA-B*07:02 | 9 | IC<sub>50</sub> | **0.75** | **0.47** | 0.51 | 0.68 | 0.47 | **0.56** |
| HLA-B*07:02 | 9 | Binary | 0.84 | 0.37 | **0.45** | **0.85** | **0.38** | 0.42 |
| HLA-B*07:02 | 9 | t1/2 | 0.78 | 0.51 | 0.39 | **0.86** | **0.59** | **0.57** |
| HLA-B*35:01 | 9 | IC<sub>50</sub> | 0.65 | **0.29** | 0.20 | **0.67** | 0.13 | **0.26** |
| HLA-B*57:01 | 9 | IC<sub>50</sub> | 0.66 | **0.27** | **0.25** | **0.70** | 0.18 | 0.21 |
| HLA-B*58:01 | 9 | IC<sub>50</sub> | **0.66** | 0.34 | **0.32** | 0.64 | **0.36** | 0.24 |
| HLA-B*57:01 | 9 | Binary | 0.82 | 0.37 | 0.40 | **0.84** | **0.39** | **0.45** |
| HLA-B*58:01 | 9 | t1/2 | **0.70** | **0.37** | 0.29 | 0.63 | 0.28 | **0.38** |
| Average |  |  | **0.72** | **0.40** | **0.32** | **0.72** | **0.40** | 0.31 |
| Count of Best |  |  | 8 | 11 | 11 | 12 | 9 | 9 |

### HONN

High-order Neural Network (HONN) is a semi Boltzmann machine based feed-forward neural network [7]. It has one hidden layer and one or more fully connected layers and used a sophiscated layer-wise training plus back-propagation based fine-tuning to train the network. In HONN, a peptide of 9 amino acids is represented by a 180-dimensional vector with each animo acid represented by its corresponding 20-dimensional substitution probabilities extracted from BLOSSUM matrix. Since there’s no code available for HONN, we trained another set of allele-specific CNN models on the BD2009 dataset and tested these models on the BLIND dataset. These two datasets were used in the HONN paper to train and evaluate their HONN model.

Table 9 shows the performance comparison between DeepMHC and HONN in terms of AUC scores. The AUC scores of HONN algorithm are extracted from the supporting material of the paper [7]. First, we found that despite HONN used additional information via BLOSOM based sequence encoding and a sophisticated training procedure, our CNN models showed competitive performance. Out of the 29 alleles test datasets, both algorithms have the same AUC scores for 13 allele datasets. Out of the remaining 16 test datasets, HONN achieved better results on 9 datasets while DeepMHC worked better over 7 datasets. Actually, out of the 9 datasets that HONN worked better, the DeepMHC’s performance scores are lower with a gap of less than 0.03 for 7 datasets. These results showed the DeepMHC achieved competitive performance without resorting to additional information via BLOSUM based peptide encoding and without the specially designed mean and covariance hidden units for modeling the mean of input features and pairwise interactions between input features. Our algorithm also is preferred in terms of much simpler computatioal training procedure.

**Table 9.** Prediction Performance of DeepMHC V.S. HONN on BLIND Dataset (AUC scores)

| Allele | DeepMHC | HONN | Allele | DeepMHC | HONN |
| --- | --- | --- | --- | --- | --- |
| A*01:01-BLIND | 0.89 | 0.89 | A*30:01-BLIND | 0.91 | 0.91 |
| A*02:01-BLIND | 0.92 | 0.92 | A*30:02-BLIND | 0.72 | **0.78** |
| A*02:02-BLIND | **0.90** | 0.86 | A*31:01-BLIND | 0.86 | **0.88** |
| A*02:03-BLIND | 0.96 | 0.96 | A*33:01-BLIND | 0.91 | 0.91 |
| A*02:06-BLIND | 0.87 | 0.87 | A*68:01-BLIND | 0.92 | 0.92 |
| A*03:01-BLIND | 0.89 | **0.92** | A*68:02-BLIND | **0.96** | 0.95 |
| A*11:01-BLIND | 0.94 | 0.94 | A*69:01-BLIND | **0.93** | 0.92 |
| A*23:01-BLIND | 0.85 | **0.86** | B*07:02-BLIND | **0.90** | 0.88 |
| A*24:02-BLIND | 0.79 | **0.81** | B*08:01-BLIND | **0.95** | 0.94 |
| A*26:01-BLIND | 0.92 | 0.92 | B*15:01-BLIND | 0.90 | **0.92** |
| A*29:02-BLIND | 0.86 | **0.89** | B*27:05-BLIND | 0.91 | 0.91 |
| B*44:02-BLIND | 0.84 | **0.91** | B*35:01-BLIND | **0.86** | 0.85 |
| B*51:01-BLIND | 0.92 | 0.92 | B*39:01-BLIND | 0.95 | **0.96** |
| B*57:01-BLIND | 0.95 | 0.95 | B*40:01-BLIND | 0.93 | 0.93 |
| B*58:01-BLIND | 0.96 | 0.96 |  |  |  |

### Comparison of Protein Sequence Encodings for CNN-based Binding Prediction

Unlike images in computer vision which can be naturally encoded as matrices, amino acid sequences need to be encoded as input vectors or matrices for training deep neural networks. The encoding scheme may have big impact on the prediction performance, which however, has not been well explored. In Section, we adopted a multi-channel encoding for all previous experimetns as illustrated in Figure 2. Here we evaluated a single-channel sequence encoding method and compared its performance to the prior approach that we used.

In multi-channel encoding, a protein sequences is encoded into a tensor object with width of 13, height of 1 and depth (channel) of 20. In the single-channel encoding approach, the tensor object can be represented as a 2D matrix. So the enocded object of a 13-amino acid sequence would have a width of 13, height of 20 and depth of 1. To compare these two encoding methods, we construct another CNN structure as showed in Figure 4. We trained and evaluated this new model (noted as DeepMHC-Row) following the same procedure described in section on allele HLA-A*02:01. The comparison results are shown in Figure 5. From the result we can see that DeepMHC with multiple-channel encoding is better than DeepMHC-ROW with single-channel encoding on all three metrics. This is expected as in one-hot encoding, each column vector corresponds to the same amino acid of the peptide, which should be encoded as multiple channels (similar to the R/G/B colors of a pixel in image recognition) instead of separate unrelated entries in a matrix.

**Figure 4.**
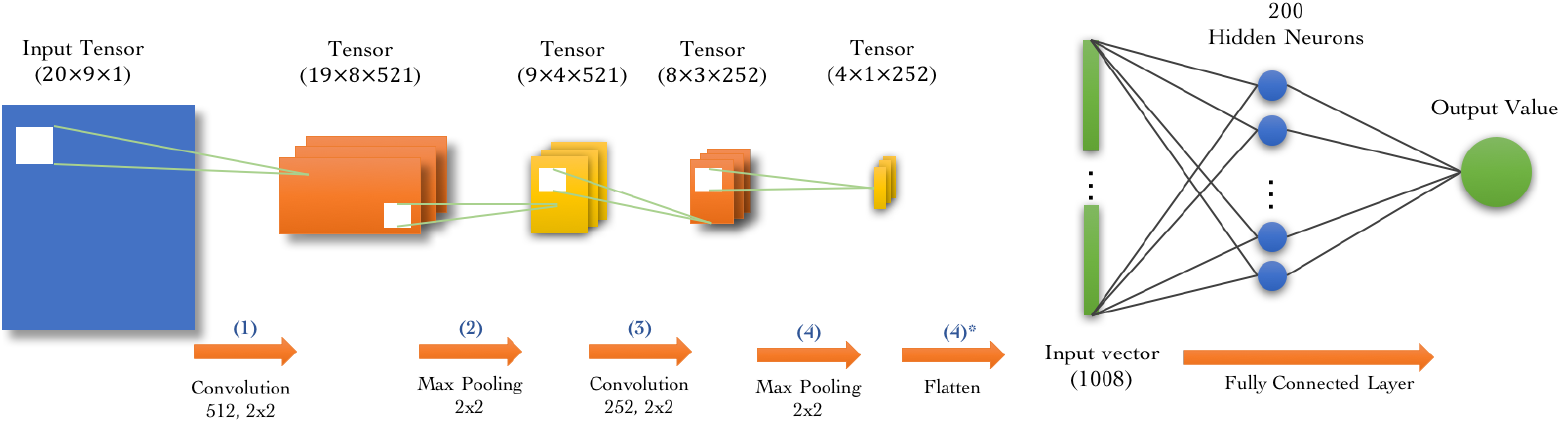
CNN Network structure of DeepMHC-Row with single-channel 2D encoding scheme. Firstly, a 9-length sequence is encoded into a 20 × 9 × 1 dimension input tensor. Each row encodes the corresponding amino acid position. Then the input tensor is fed to the (1) convolution layer which has 512 filters with 2 × 2 receptive field size. Next in layer (2), output tensor of (1) is applied with a Max-Pooling layer with a 2 × 2 window. After Max-Pooling, another convolution layer (3) with 252 filters with 2 × 2 size is applied to the output of the pooling layer. Then another) Max-Pooling is applied on the tensor. A flatting operation is applied on the layer (4)’s output and a 1D vector with 1008 values is obtained, which serves as the input of the fully connected layer (4), which has 2 hidden neurons. Finally, layer (4) gives the predicted value.

**Figure 5.**
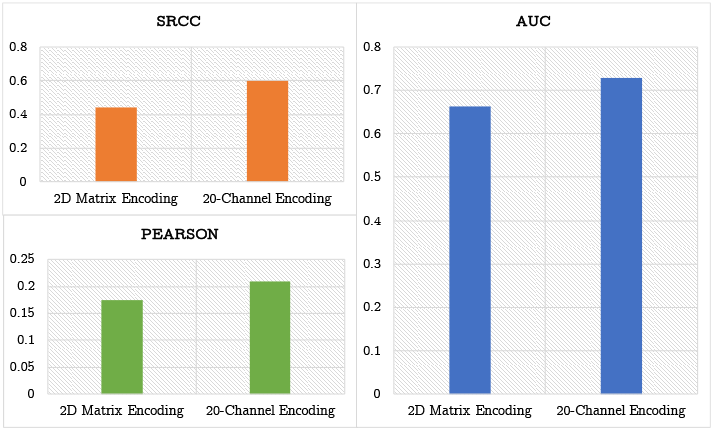
Prediction performance of 20-channel encoding vs 1-channel encoding. Trained and tested two models on allele HLA-A*02:01.

### Exploring Hyperparameters of CNN models for Binding Prediction

In addition to protein sequene encoding, there are several hyperparameters of the CNN that need to specify appropriately to achieve good prediction performance. Here we explored different network structures by tuning the network hyperparameters including: the size and number of convolution filters, the depth of networks, and the batch size of training samples. The optimal result is what we used in the final CNN network structure in Figure 2.

First, we evaluated how the number of filters of the convolutional layer affects the prediction performance. We used the CNN model shown in Figure 2 but varied the number of filters of the convolution layer by setting it as 25, 50, 100, 250, and 512 and evaluated the prediction performance of the resulting CNN models on 4 allele datasets: HLA-A*02:01, HLA-A*68:01, HLA-A*02:06 and HLA-B*58:01. We trained these prediction models on BD2013 and tested them on the weekly benchmark data in the same way as we did in Section.

Figure 6 shows the prediction performance of the CNN models over the alle HLA-A*02:01 dataset with different numbers of filters on the four datasets. First we found that when the number of filters is 25, the performances are all very low in terms of AUC/SRCC/Pearson compared to larger number of filters. When the filter numer increases to 50 and up to 250, the performances also improved though the performances of CNNs with 50/100/250 filters did not vary a lot. However, when the number of filters increased to 512, the performances in terms of all three criteria all jumped significantly. In all previous CNN models for DNA-binding [11] and peptide binding prediction algorithms [15, 16], the number of filters are all set to a few dozens from 30 to 100. The demand for a larger number of convolution filters in peptide binding prediction compared to DNA/RNA binding prediction is probably due to the fact that the 20-channel encoding of amino acids leads to a significantly larger encoding space, which needs a large number of convolution filters. The reason is that in convolutional neural networks, the number of convolution fitlers of the convolution layer determines the number of diverse building block patterns that they need to model for higher level prediction. Each filter is supposed to capture a basic feature representation of input tensors. While a few dozens of filters such as 30-60 filters are sufficient to achieve good performance in DNA binding prediction [11], it is not the case for the peptide-binding prediction since here the alpahbet size of amino acids is now 20 instead of 4 of nucleotides. As shown in our one-hot encoding, the 20-D encoding for each amino acid and the 9 or 10 amino acids lead to a much diverse pattern space, thus requiring much more convolution filters to learn to extract these patterns. This may explain why our DeepMHC can achieve much better performances than those CNN models in [15, 16], both using about 30-60 filters. In our case, the input tensor has a dimension of 1 × 13 × 20 and 512 filters are used in the CNN models, whose performance is demonstrated by the competitive results across all datasets using a simple one-hot encoding scheme.

**Figure 6.**
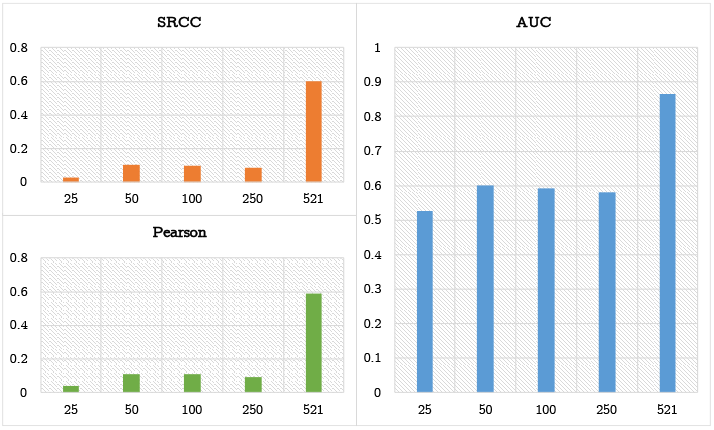
Influence of the number of filters on prediction performance. We compared the performance of DeepMHC with different numbers of convolution filters on the dataset of alle HLA-A*02:01. We found that the CNN model with 512 fitlers achieved the best results in terms of three performance metrics.

Another hyperparameter of our CNN model is *batch size*, which is the number of samples fed to the neural network each time for calculating the errors and updating the weights of the neural network in the training stage. Compared to sample feeding one by one, batch training allows to take advantage of the fast array operations and hence faster training time. However, too large batch size may lead to overfitting.

To explore how the batch size influences the prediction performance, we trained CNN models using full-batch (or batch) and different sizes of mini-batches for the HLA-A*02:01 allele dataset. The models were then tested on the IEDB weekly dataset. Figure 7 shows the prediction performances over the HLA-A*02:01 dataset as evaluated by AUC, SRCC, and Pearson coefficient. First we found that the full-batch training got the worst results in terms of all three evaluation criteria. In terms of Pearson coefficient, the batch size 250 achieved the best result. In terms of AUC and SRCC, the performances do not vary too much except for the full-batch training. Theoretically, the full-batch training should lead the model to the best optimization direction during SGD training. A possible reason for its low performance is that we used the same learning rate in all training processes. Wilson et al. [30] suggested that when dealing with full batch training, a much smaller learning rate is needed and a full-batch training estimates the gradient only at the starting point in the weight space, and thus cannot follow curves in the error surface.

**Figure 7.**
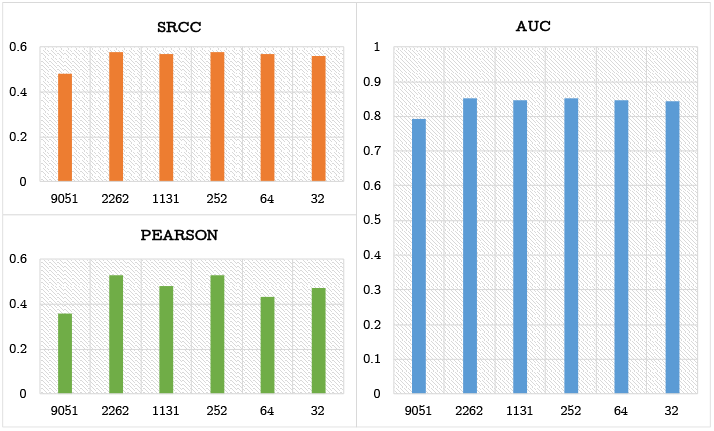
Influence of batch size on prediction performance over the allele HLA-A*02:01 dataset. Testing models are trained with different batch sizes: full batch (all 9051 training samples), 1/4 of full batch (2262 samples), 1/8 of full batch (1131 samples), 252 samples, 64 samples, 32 samples. The results shown are over the binary data. Similar performance trends have been observed on resutls for the IC50 dataset and the 11/2 dataset.

The last hyperparameter that we explored is the number of convolution layers. In our final CNN model shown in Figure 2, two convolution layers and one fully connected layer are used, which maybe considered as “not so deep”. However, this structure choice is based on our evaluation of the performances of deeper netowrks as Figure 8 shows.

**Figure 8.**
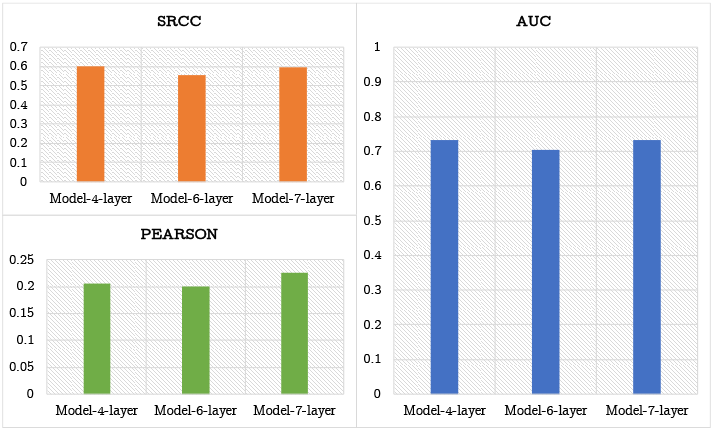
Influence of CNN network depth on the prediction performance. Tested with 4-layer model, 6-layer model and 7-layer model on allele Allele HLA-A*02:0?s data.

From the performance results in Figure 8, no significant differences were observed for CNN models with deeper layers including 4-layer(conv-pool-conv-full), 6-layer (conv-pool-conv-pool-conv-full) and 7-layer(conv-pool-conv-pool-conv-pool-full). A possible reason is that the dimension of our input tensor is 1 × 13 × 20. Usually, a convolution layer in above models uses kernels of size 1 × 2 or 1 × 3. After too many layers, the intermediate tensors will be very short. So after several layers, it loses the power to serve as more abstract high-level features.

### Transfer Learning on MHC Alleles with Small Number of Samples

A major issue in MHC-I peptide binding prediction is that some of alleles only have a very limited number of binding peptide samples, which makes it challenging to train accurate prediction models. An effective approach to address this issue is to use transfer learning[31], which improves a learner from one domain by transferring information from a related domain. We would like to see if transfer learning can improve the prediction performance for MHC-I alleles with small number of traing samples by fine-tuning the pretrained models trained with samples of another allele with more samples. To verify this approach, we picked two datasets: allele HLA-A*02:01 and allele HLA-A*02:02. Both are clustered into same supertype A02, one of 6 supertypes that the whole HLA-A alleles are clustered into [32]. The HLA-A*02:01 dataset has the most training samples (9051) and HLA-A*02:02 has 2465 training samples. We first train 5 CNN models using the samples from the allele HLA-A*02:01 dataset. And then we trained 5-fold cross-validation models for HLA-A*02:02 following the same procedure described in Section except that before training each fold’s model, we initialized the network’s weights of the first convolution layer as those of one of the 5 pretrained models of allele HLA-A*02:01. Here the transfer learning is conducted by initializing weights learned from the related dataset allele HLA-A*02:01.

Figure 9 shows the performance improvement by transfer learning in terms of three performance criteria. We found that the model trained with pretrained weights has achieved a much better prediction performance. The AUC has increased to 0.79 from 0.73; the SRCC reaches 0.72 from 0.66; and the Pearson changes from −0.03 to 0.01. When comparing this improved results with those reported in Table 4, this transfer learning based model achieved the best results on test dataset HLA-A*02:02(IC5o). Such significant performance improvement by transfer learning may be partially due to the fact that we were actually doing a “good” initialization of weights for the first convolutiona layers by reusing the weights from the pretrained CNN models of HLA-A*02:01. It is increasingly recongized that proper initiallization of the weights of the convolution layers is helpful for the CNN model to get better prediction performance [33, 34, 35].

**Figure 9.**
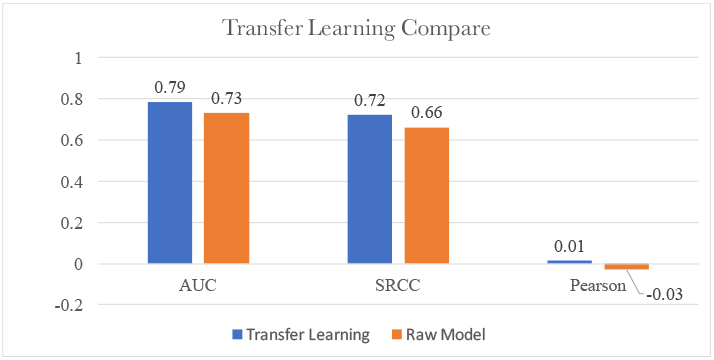
Performance improvement by transfer learning on allele HLA-A*02:02 dataset using pretrained Models from allele HLA-A*02:01. The transfer learning model and the raw model were both evaluated on the same weekly benchmark dataset from IEDB.

## Discussion

Convolutional neural networks are powerful prediction models that have demonstrated their potential in DNA/RNA binding predictions. The hierarchical feature extraction and end-to-end learning allow them to capture the non-linear relationships among the amino acid positions of the peptides as regard to the binding affinity. Our extensive experiments showed that CNN models can outperform all existing non-deep learning MHC-I binding predictors (such as the complicated NetMHCpan) using only the amino acid sequences alone.

However, design of a convolutional neural network for protein-peptide binding prediction needs to identify appropriate settings and parameters to match the target task. For example, our experiments found that the number of convolution filters must be much larger than those used in DNA-binding predictors due to the significantly larger search space. Due to the limited number samples for each allele dataset, it is found that increasing the number of convolution/max-pooling layers more than 3 layers does not necessarily improve the prediction performance. We also found that the multi-channel one-hot encoding works much better than the naive 2D matrix encoding. These extensive parameter exploration experiments help to design better CNN models for MHC-I peptide binding predictors.

In our current DeepMHC model, we only used the amino acid sequences without any additional structural, physichochemical, or evolutionary information and yet our algorithm is competitive and most of time better than all those algorithms that use these information. We also demonstrated that CNN models can achieve better or competitive results than other deep learning and CNN models without using the sophisticated word embeeding representation or the BLOSOM information. This does not mean that our model cannot take advantage of those additional information. How to incorporate those addtional information effectively into our CNN framework is a topic to be investigated.

Another major issue in MHC-I peptide binding prediction is the limited number of sample for some alleles. This is especially for deep learning approaches, which usually needs a large number of samples to perform well. However, our preliminary exploration of using all samples of different alleles to train a single Pan-specific model failed. One idea is to incorporate the receptor sequence into the model together with the peptide sequences as NetMHCSpan does.

## Conclusion

We have developed DeepMHC, a deep convolutional neural network based peptide binding prediction algorithm, which is featured with unified models for all allele MHC-I molecules and simple one-hot encoding. Extensive experiments on MHC-I peptide binding prediction showed that DeepMHC achieved state-of-the-art performance over a majority of the benchmark datasets from IEDB. It is found that the success of our deep CNN model with one-hot encoding is based on the automated learning of non-linear high-order dependency among amino positions on the peptide. We found that a large number of filters (¿500) with small filter size is essential for its high performance. Our approach can be readily applied to other protein-peptide binding prediction problems by only replacing the training data sets without much hyper-parameter messing. The open-source source code of DeepMHC will be availalbe at http://mleg.cse.sc.edu once the paper is accepted.

## Acknowledgments

We gratefully acknowledge the support of NVIDIA Corporation with the donation of the K40 Pascal GPU used for this research.

